# Streaming algorithms for identification of pathogens and antibiotic resistance potential from real-time MinION^TM^ sequencing

**DOI:** 10.1101/019356

**Authors:** Minh Duc Cao, Devika Ganesamoorthy, Alysha G. Elliott, Huihui Zhang, Matthew A. Cooper, Lachlan Coin

## Abstract

The recently introduced Oxford Nanopore MinION platform generates DNA sequence data in real-time. This opens immense potential to shorten the sample-to-results time and is likely to lead to enormous benefits in rapid diagnosis of bacterial infection and identification of drug resistance. However, there are very few tools available for streaming analysis of real-time sequencing data. Here, we present a framework for streaming analysis of MinION real-time sequence data, together with probabilistic streaming algorithms for species typing, multi-locus strain typing, gene presence strain-typing and antibiotic resistance profile identification. Using three culture isolate samples as well as a mixed-species sample, we demonstrate that bacterial species and strain information can be obtained within 30 minutes of sequencing and using about 500 reads, initial drug-resistance profiles within two hours, and complete resistance profiles within 10 hours. Multi-locus strain typing required more than 15x coverage to generate confident assignments, whereas gene-presence typing could detect the presence of a known strain with 0.5x coverage. We also show that our pipeline can process over 100 times more data than the current throughput of the MinION on a desktop computer.

## Background

Massively parallel, short-read sequencing has profoundly transformed genomics research [1, 2] and has become the dominant technology for sequencing DNA. 5 However, one inherent limitation of sequencing millions of sequence fragments in parallel one base at a time is that the sequencing run has to finish before the data analysis can begin. As a result, sequence analysis algorithms have been designed to make inference on 10 a complete sequencing dataset. In contrast, streaming algorithms are a class of algorithms which are applied to a sequence of data events and typically maintain an internal summary of the data as well as an approximation to the full inference without needing to store all of 15 the observations [3]. Streaming algorithms have application in particle and solar physics, computer network analysis and finance [4].

Oxford Nanopore Technologies has recently released a portable MinION sequencing device, which utilizes 20 the nanopore sequencing technology proposed in the 1990s [5]. The key innovation of this device is that it measures the changes in electrical current as a singlestranded DNA passes through the nanopore and uses the signal to determine the nucleotide sequence of the DNA strand [6, 7]. This sequence data can be retrieved and analyzed as it is generated, providing the opportunity to obtain answers in the shortest possible time. Real-time sequencing has immense potential in many applications, especially in time-critical areas such as rapid clinical diagnosis.

In order to realise this potential there is a need to develop streaming bioinformatics algorithms which continually update inference about the sample as each sequence read is generated. To be of practical use – for example to know when to when to make a diagnosis in the clinic - these algorithms must continuously update not only a point estimate (e.g. which species present and their proportions), but also confidence intervals in that estimate. Several systems incorporating real-time analysis of MinION data have been developed recently such as the cloud based platform Metrichor (Oxford Nanopore), work by Quick et al [8] and MetaPORE [9], focusing on placing the sample on a phylogenetic tree but without providing an estimate of the confidence in this assignment.

Here we present a flexible framework for real-time analysis on MinlON sequence data directly as it is sequenced and base-called. The framework can incorporate multiple real-time analyses to suit the problems at hand and can be deployed on a single computer or on a high performance computing facility and computing cloud. We also present four streaming algorithms for identification and characterization of pathogen samples. These algorithms, which are seamlessly integrated into the pipeline, report analysis results along with their confidence levels so that users can decide when to stop a sequencing run.

By sequencing three bacterial isolate samples and a mixture sample on the MinION sequencer, we demonstrate that we can reliably determine the species and strain type of a sequenced sample with only 500 reads. This was achieved in less than half an hour of sequencing with the current throughput of the MinION. Furthermore, we show that we can identify the majority of the drug resistance genes present in a sample within 2 hours of sequencing, and the full drug resistance profile within 10 hours. We also show that MinION sequence data can be used for accurate Multi-Locus Sequence Typing (MLST), despite the relatively high error rates associated with the technology. The pipeline can perform all these analyses on a single computer at a throughput of over 100 times higher than our best runs. As the throughput of nanopore sequencing is expected to increase, the time to obtain these results will be significantly shortened. Our findings support the potential use of MinION sequencing for real-time analysis of clinical samples for species detection and analysis of antibiotic resistance.

## Results and discussion

### Real-time analysis framework

Our real-time analysis framework consists a number of *streaming* programs communicating to each other via the network sockets or the inter-process communication pipes provided by Unix-like operating systems. These programs typically take a sequence of items as input and process after every some small number of items arrive. They either retain only the relevant statistics of the data, or upon processing any data items, immediately forward only the necessary information to the downstream programs for further processing. Processing data in this way requires a much smaller memory footprint and hence is relevant for processing large amount of data, especially real-time data from MinION sequencing.

We developed a number of auxiliary programs to facilitate setting up a real-time pipeline for analysis of MinION sequencing data. These include scripts for setting up communication channels in a pipeline, thereby allowing the pipeline to be deployed on a high performance computing cluster to scale with massive amounts of data. Programs for simple analyses of the MinION sequencing data such as initial analysis (npReader [10]) and read-filtering on the basis of read length and read quality are also provided.

We developed streaming algorithms for a handful of identification problems, namely species typing, multi-locus strain typing; gene-presence strain-typing and identification of antibiotic resistance profiles (see Methods). We integrated the implementations of these algorithms into the analysis pipeline (see Figure 1). In this pipeline, npReader [10] continuously scans the folder containing sequencing data in parallel with the MinION sequencing. It picks up sequenced reads as soon as they are generated (from Metrichor), and simultaneously streams them through the pipeline for the identification analyses. The pipeline also makes use of off-the-shelf bioinformatics tools such as BWAMEM [11] as described later. In each step of this pipeline, data is piped from one process to the next without being written to disk, with the exception of base-calling via Metrichor in which each read is written to disk once it has been base-called, and then read almost immediately by npReader.

**Figure 1.**
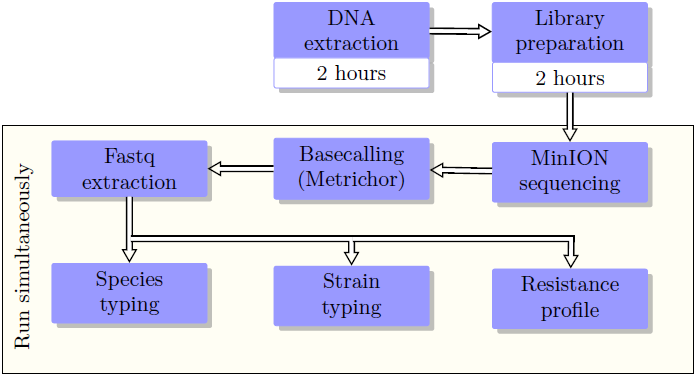
Schematic of the real-time analysis pipeline. Typically it takes at least 4 hours to go from a cultured isolate to a library for MinION sequencing. Once the MinION starts sequencing, DNA fragments are sequenced (on the MinION) and base-called (by Metrichor cloud) instantaneously, and are simultaneously streamed through the pipeline. Analysis results and their confidence levels are reported in real-time. User can stop an analysis or the whole pipeline once the desired confidence levels are obtained.

We evaluated our real-time analysis pipeline and the accuracy of our algorithms using five MinION sequencing data sets. Four of these datasets were collected before the pipeline was developed, and hence we emulated the timing of the sequencing for the evaluation from these datasets. Specifically, we extracted the time that each read was sequenced, and streamed the sequence reads in the exact order and timing into the pipeline. With the emulation, we were able to stream the sequencing data with a hypothetical throughput of 120 times higher what we could obtain with the MinION. This allowed us to test the scalability of the pipeline against the projected future throughput such as from the PromethION platform. The fifth dataset was passed through our pipeline as it was base-called from Metrichor, and thus represents a true demonstration of the real-time capability of the pipeline. Finally, we validated the analysis results by sequencing these samples with Illumina MiSeq platform, where bioinformatics analysis methods are well-established.

### Data generation

We prepared two samples of cultured isolates of the *Klebsiella pneumoniae* strains ATCC BAA-2146, ATCC 13883; one *Klebsiella quasi-pneumoniae* strain ATCC 700603 and a library mixture sample. This mixture sample contains two different sequencing libraries prepared from the *Escherichia coli* strain ATCC 25922 and the *Staphylococcus aureus* strain ATCC 25923, pooled at different levels prior to sequencing (Table 1). We sequenced sample ATCC BAA-2146 and ATCC 700603 with the MinION using chemistry R7, the others using the improved chemistry R7.3 (see Methods).

**Table 1.**
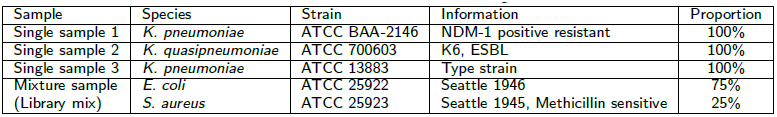
Details of the four samples.

| Sample | Species | Strain | Information | Proportion |
| --- | --- | --- | --- | --- |
| Single sample 1 | *K. pneumoniae* | ATCC BAA-2146 | NDM-1 positive resistant | 100% |
| Single sample 2 | *K. quasipneumoniae* | ATCC 700603 | K6, ESBL | 100% |
| Single sample 3 | *K. pneumoniae* | ATCC 13883 | Type strain | 100% |
| Mixture sample<br>(Library mix) | *E. coli* | ATCC 25922 | Seattle 1946 | 75% |
|  | *S. aureus* | ATCC 25923 | Seattle 1945, Methicillin sensitive | 25% |

In order to validate the analysis results from MinION sequencing, we sequenced all aforementioned isolates with the established Illumina platform MiSeq to a coverage exceeding 100-fold. Isolates in the mixture sample were sequenced separately. We assembled the MiSeq sequencing reads to obtain high quality assemblies of the five strains. With the assemblies, we were able to identify the strain types and the antibiotic resistance profiles of these strains (see Methods). These results were used as the benchmarks to validate the analysis of MinION sequencing data.

### Sequencing yields and quality of MinION sequencing

Sequence reads from the MinION were classified into three types: *template, complement* and higher quality *2D* reads (*i.e*., reads resulted from computationally merging a template and a complement read). The average Phred quality of template and complement reads across four runs was in the region of 5 while 2D reads were in higher quality, with average Phred quality about 9 (see Table 2 and Figure 3). The median read lengths of three *K. pneumoniae* samples were approximately 5Kb, while the mixture sample was only less than 1Kb (Figure 3). We observed a variation in terms of sequence yields across the four runs. While we obtained nearly 40000 reads (185Mb) for sample *K. pneumoniae* ATCC BAA-2146 after 60 hours of sequencing, the run for sample *K. quasipneumoniae* ATCC 700603 yielded only 7092 reads (39Mb) with the same running time (Figure 2). We sequenced sample *K. pneumoniae* ATCC 13883 and the mixture sample for 36 and 20 hours respectively both with the chemistry 7.3 but and the yields were markedly different to each other. The read length and accuracy of our runs were consistent with other user reports [12–15].

**Table 2.**
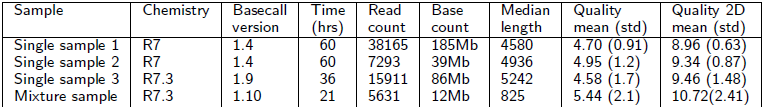
Details of the four MinION sequencing runs.

**Figure 2.**
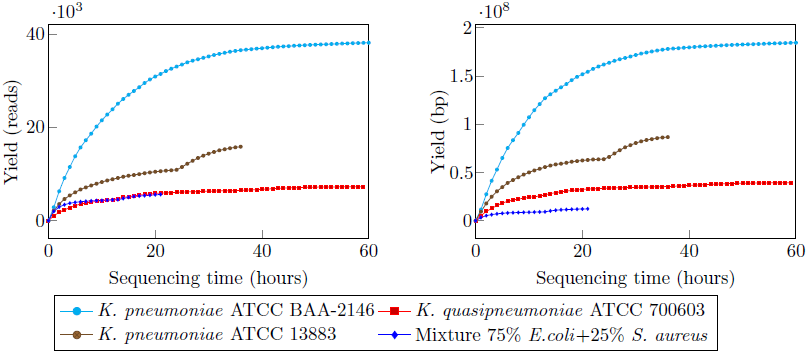
Sequencing yields over time for the four samples. Yields are shown in terms of read count (left) and base count (right).

**Figure 3.**
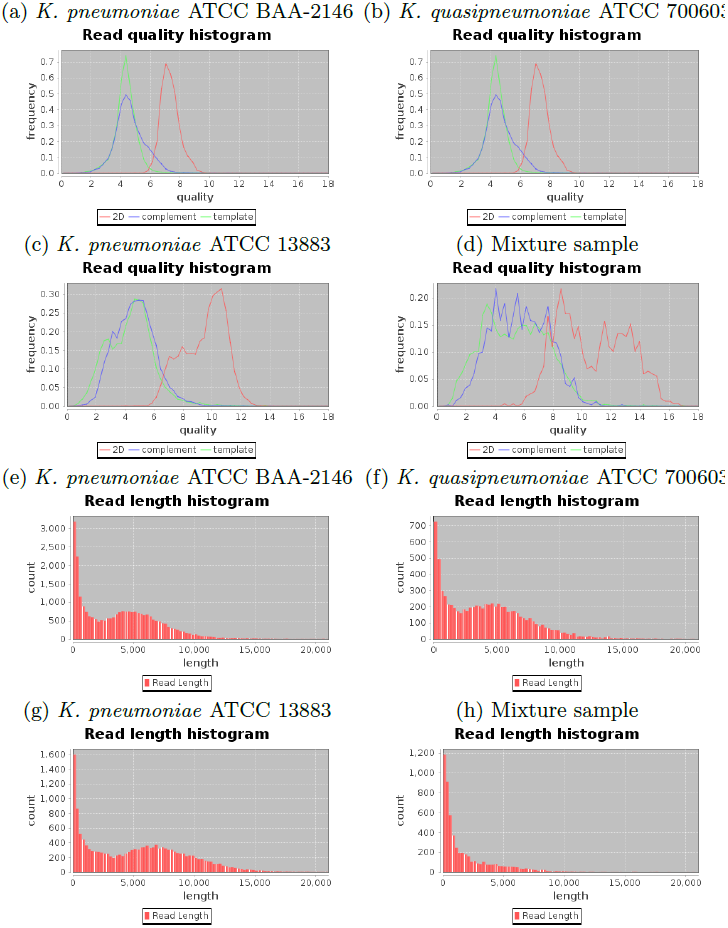
The distribution of read quality and read length of the four MinION runs.

### Species detection

For real-time bacterial species detection, we built a database from 2,785 complete genomes of 1,489 bacterial species available in GenBank (

~~~
http://www.ncbi.nlm.nih.gov/genbank/
~~~

, accessed Nov 2014), augmented with two *K. quasipneumoniae* genomes (which was not the strain we sequenced) as none were present in the database. The database contained a number of *K. pneumoniae, E. coli* and *S. aureus* strains (10, 63 and 49 respectively), but none of the five strains in our samples were present. The pipeline aligns sequence reads as they are generated from the sequencer to this database. The species typing algorithm periodically computes the simultaneous proportions of the species present in the sample and reports the 95% confidence intervals of these proportions (See Methods).

In both *K. pneumoniae* samples as well as the *K. quasipneumoniae* sample, we successfully detected the major species present in the isolate. This was achieved with as little as 120 sequence reads requiring only 5 minutes of sequencing time (Figures 4a), b) and c)). For *K. pneumoniae* strains ATCC BAA-2146 and ATCC 13883, it required less than 500 reads (10 and 15 minutes of sequencing, respectively) to reach a 95% confidence interval of less then 0.05. For strain ATCC 700603 it required only 300 reads to correctly identify *K. quasipneumoniae* as the species.

**Figure 4.**
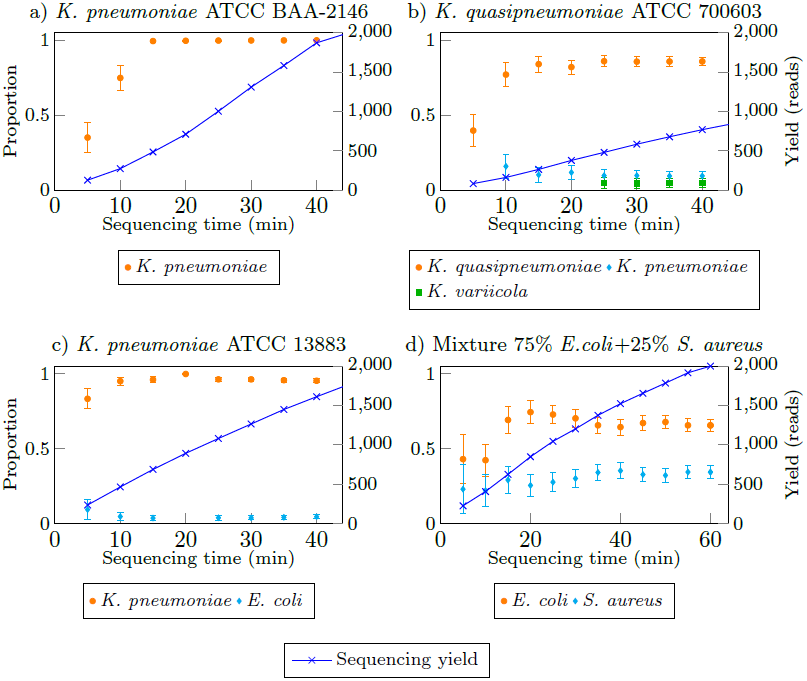
Real-time identification of bacterial species from MinlON sequencing data for four different bacterial samples. The bars represent confidence intervals at 95% level.

The pipeline accurately identified the two species in the mixture sample as *E. coli* and *S. aureus* after obtaining around 100 reads (5 minutes of sequencing). The reported proportions became stable after around 1200 reads (35 minutes of sequencing). *E. coli* was the predominant species type in the mixture sample and it was evident with high proportion of sequencing reads supporting the *E. coli* species.

### Multi-locus Sequence Typing

*K. pneumoniae* and other bacteria are conventionally strain typed using a MLST system which requires accurate genotyping to distinguish the alleles of seven house-keeping genes [16]. Our analysis of MinlON raw read quality (Fig. 3), together with other user reports [12–15], indicated high error rates in MinlON sequencing in comparison to Illumina Miseq sequencing. This suggested that MLST typing was challenging with MinION sequence data, especially in real-time fashion.

We developed a method to carry out MLST typing using MinION sequence data. Our method selected only reads spanning one of the house-keeping genes. It then used multiple reads aligned to the same gene to correct error in the raw sequence reads and subsequently combined information across multiple alleles in a likelihood-based framework (see Methods). Table 3 presents the top five highest score types (in log-likelihood) forK. *pneumoniae* and *K. quasipneumoniae* strains using MinION sequencing. In all three strains, the correct types were the highest score out of 1678 types available in the MLST database. We noticed that the typing system also outputted several other strain types with the same likelihood (*e.g.*, types ST-751 and ST-864 for strain ATCC BAA-2146 and type ST-851 for strain ATCC 700603). We examined the profiles of these types, and found that these strain types were highly similar. For example, strain types ST-751 and ST-864 (reported for strain ATCC BAA-2146) differed to the correct strain type ST-11 by only one SNP from the total of 3012 bases in seven genes. Similarly, strain type ST-851 (co-highest score reported for strain ATCC 700603) differed to the correct strain type ST-489 by two alleles (genes phoE and tonB). There was only one read aligned to these two genes by the end of the run due to the poor yield of this run, which may have also contributed to inability to differentiate these two strain types. While the results were encouraging, this also suggested that a more accurate strain-typing methodology would need to consider all of the sequenced reads, rather than just those covering 7 house-keeping genes. Therefore we further devised a method for strain-typing which was based on presence or absence of genes.

**Table 3.** Multi-locus strain-typing results for three *K. pneumoniae* strains. The top five probable MLST types are shown for each sample. The highest score strain types are **highlighted**.

| ATCC BAA-2146<br>ST-11 |  |  | ATCC 700603<br>ST-489 |  | ATCC 13883<br>ST-3 |  |
| --- | --- | --- | --- | --- | --- | --- |
| Rank | Type | Score | Type | Score | Type | Score |
| 1 | **ST-11** | **1985.47** | **ST-489** | **418.45** | **ST-3** | **1451.65** |
| 2 | ST-751 | 1985.47 | ST-851 | 418.45 | ST-136 | 1450.21 |
| 3 | **ST-864** | **1985.47** | ST-257 | 413.57 | ST-38 | 1444.81 |
| 4 | ST-1080 | 1984.46 | ST-356 | 413.57 | ST-1106 | 1444.19 |
| 5 | ST-1680 | 1982.62 | ST-414 | 413.57 | ST-931 | 1441.44 |

### Strain typing by presence or absence of genes

We developed a novel strain typing method to identify the bacterial strain from the MinION sequence reads based on patterns of gene presence and absence. We downloaded the genome assemblies of all strains for *K. pneumoniae, E. coli* and *S. aureus* species from RefSeq repository and identified their strain types using the relevant MLST schemes. This resulted in sets of 125 strain types for *K. pneumoniae*, 353 for *E. coli* and 107 for *S. aureus*. For each strain type, we picked the highest quality assembly (in terms of N50 statistic) and extracted gene sequences from its RefSeq gene annotation. We then grouped genes from a species based on 90% sequence identity, and therein obtained the gene profile for each strain type.

Our pipeline identified genes present in the sample from sequence reads as they were generated by the MinION device. It then used this information to infer the posterior probability of each of the strain types, as well as the 95% confidence intervals in this estimate (see Methods). For our *K. pneumoniae* and *K. quasipneumoniae* samples, we successfully identified the corresponding strain types from the sequence data with 95% confidence within 10 minutes of sequencing time and with as few as 200 sequencing reads (Figures 5a., b., and c.). We streamed sequence reads from the mixture sample through the strain typing systems for *E. coli* and *S. aureus*, and in both cases, the correct strain types of two species in the sample were also recovered. The correct type for *E. coli* strain in the 75%/ 25% *E. coli,S. aureus* mixture was recovered after 25 minutes of sequencing with about 1000 total reads (or approximately 750 *E. coli* derived reads). (Figure 5d.). The pipeline was able to correctly predict the *S. aureus* strain (which is known to have much less gene content variation) in this mixture sample after two hours of sequencing with about 2,800 total reads (or approximately 700 *S. aureus* derived reads).

**Figure 5.**
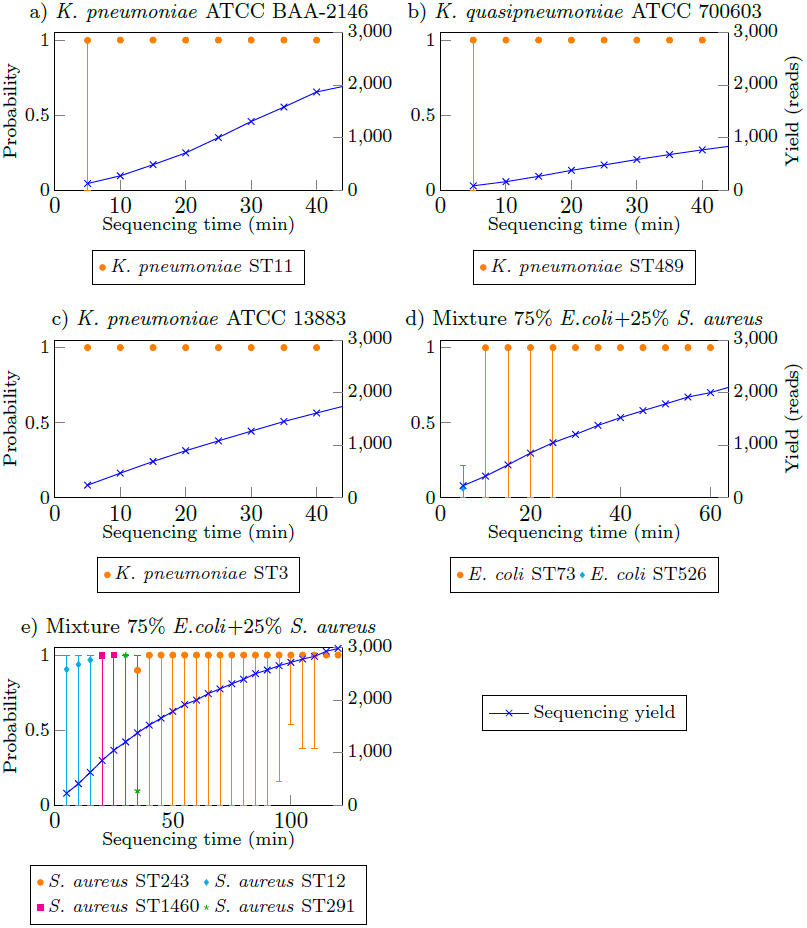
Real-time identification of strain type from MinION sequencing data on three different *K. pneumoniae* strains and a mixture sample of an *E. coli* and a *S. aureus* strain. The bars represent confidence intervals at 95% level.

### Antibiotic resistance detection

The antibiotic resistance gene profiles of the samples were also characterized with MinION sequencing data. We obtained antibiotic drug resistance genes from *ResFinder* database [17] (

~~~
https://cge.cbs.dtu.dk/services/ResFinder/
~~~

, accessed July 2015). This set contained over 2132 gene sequences, including variants of the same genes. We grouped these gene sequences based on 90% sequence identity into 609 groups. In this grouping, we found that sequences in a group were variants of the same gene.

Our antibiotic resistance profile identification pipeline aligned sequence reads to this antibiotic gene database. The algorithms retained reads that aligned to these genes, and periodically performed multiple alignment of reads that were aligned to the same gene. It then generated a consensus sequence from these reads, and used a probabilistic Finite State Machine [18] to realign the consensus sequence to the gene sequence (see Methods). The pipeline reported the presence of a resistance gene as soon as the alignment score reached a threshold.

Table 4 shows the time-line of antibiotic genes detection from MinION sequencing of three *K. pneumoniae* strains. For the NDM-1 producing strain ATCC BAA-2146, we identified the presence of 26 antibiotic resistance genes in the MiSeq assembly of the strain. Our real-time pipeline identified all these 26 genes and an additional gene blaSHV from 10 hours of MinION sequencing. No further gene was detected thereafter. As gene blaSHV was reported with high confidence from the our real-time analysis, we further investigated the alignment of the MiSeq assembly with this gene, and found that the gene was actually aligned to two contigs in the assembly suggesting the MiSeq assembly might have been fragmented in the middle of the gene. We sourced a high quality assembly of the strain’s genome using PacBio sequencing [19] and found that the assembly actually contained the gene. In other words, our pipeline detected precisely the antibiotic gene profile for this strain from 10 hours of MinION sequencing. We observed that the majority of these genes were identified in early stage of sequencing, *i.e.*, three quarters of these genes were reported within 1.5 hours of sequencing, at less than 4000 reads (making up only a 3-fold coverage of the genome). We observed similar performance for *K. pneumoniae* strain ATCC 13883 where 5 out of 6 genes after two hours of sequencing. The last gene (oqxB) was detected after 9.5 hours of sequencing, again recovering the full resistance profile without any false positive. For the multi-drug resistant *K. quasipneumoniae* strain ATCC 700603, the pipeline only detected 8 out of 11 genes. The reduced sensitivity for this sample was most likely due to the low sequence yield (33Mb of data in total, or only 7-fold coverage of the genome).

**Table 4.** Timeline of resistance genes detection from the *K. pneumoniae* samples. TP/FP: true positives/false positives according to the resistance gene profiles obtained from MiSeq sequencing. *Gene blaSHV was detected from MinlON sequencing of *K. pneumoniae* ATCC BAA-2146 but not from MiSeq sequencing due to the inability to resolve a repeat in the gene.

| Time (mins) | genes | Class | TP/FP | Sensitivity (%) | Specificity (%) | Data |
| --- | --- | --- | --- | --- | --- | --- |
| *K. pneumoniae* ATCC BAA-2146 |  |  |  |  |  |  |
| 30 mins |  |  |  |  |  | 1228 reads |
|  | mphA | macrolide | TP |  |  |  |
|  | blaSHV | beta-lactamase | FP* |  |  |  |
|  | strA | aminoglycoside | TP |  |  |  |
|  | blaTEM | beta-lactamase | TP |  |  |  |
|  | strB | aminoglycoside | TP |  |  |  |
|  | blaCTX | beta-lactamase | TP | 26.67 | 87.50 |  |
| 60 mins |  |  |  |  |  | 2613 reads |
|  | blaLEN | beta-lactamase | TP |  |  |  |
|  | sul2 | sulphonamide | TP |  |  |  |
|  | blaOXA | beta-lactamase | TP |  |  |  |
|  | aac3 | aminoglycoside | TP |  |  |  |
|  | aac6 | aminoglycoside | TP |  |  |  |
|  | blaCMY | beta-lactamase | TP |  |  |  |
|  | blaCFE | beta-lactamase | TP |  |  |  |
|  | blaLAT | beta-lactamase | TP |  |  |  |
|  | blaBIL | beta-lactamase | TP | 53.33 | 94.12 |  |
| 90 mins |  |  |  |  |  | 3844 reads |
|  | QnrB | quinolone | TP |  |  |  |
|  | aadA | aminoglycoside | TP |  |  |  |
|  | oqxA | quinolone | TP |  |  |  |
|  | tetA | tetracycline | TP |  |  |  |
|  | oqxB | quinolone | TP | 76.67 | 95.83 |  |
| 120 mins |  |  |  |  |  | 5258 reads |
|  | dfrA | trimethoprim | TP | 80.00 | 96.00 |  |
| 240 mins |  |  |  |  |  | 10788 reads |
|  | blaOKP | beta-lactamase | TP | 83.33 | 96.15 |  |
| 270 mins |  |  |  |  |  | 11931 reads |
|  | rmtC | aminoglycoside | TP | 86.67 | 96.43 |  |
| 300 mins |  |  |  |  |  | 13022 reads |
|  | sul1 | sulphonamide | TP |  |  |  |
|  | sul3 | sulphonamide | TP | 93.33 | 96.55 |  |
| 540 mins |  |  |  |  |  | 20200 reads |
|  | fosA | fosfomycin | TP | 96.67 | 96.67 |  |
| 600mins |  |  |  |  |  | 21546 reads |
|  | blaNDM | beta-lactamase | TP | 100.00 | 96.77 |  |
| *K. quasipneumoniae* ATCC 700603 |  |  |  |  |  |  |
| 30 mins |  |  |  |  |  | 582 reads |
|  | oqxA | quinolone | TP |  |  |  |
|  | blaSHV | beta-lactamase | TP |  |  |  |
|  | oqxB | quinolone | TP | 27.27 | 100.00 |  |
| 60 mins |  |  |  |  |  | 1090 reads |
|  | aadB | aminoglycoside | TP | 36.36 | 100.00 |  |
| 390 mins |  |  |  |  |  | 3704 reads |
|  | sul1 | sulphonamide | TP |  |  |  |
|  | sul3 | sulphonamide | TP | 54.55 | 100.00 |  |
| 420 mins |  |  |  |  |  | 3810 reads |
|  | blaOXA | beta-lactamase | TP | 63.64 | 100.00 |  |
| 540 mins |  |  |  |  |  | 4156 reads |
|  | blaOKP | beta-lactamase | TP | 72.73 | 100.00 |  |
| *K. pneumoniae* ATCC 13883 |  |  |  |  |  |  |
| 30 mins |  |  |  |  |  | 1264 reads |
|  | fosA | fosfomycin | TP | 16.67 | 100.00 |  |
| 60 mins |  |  |  |  |  | 2186 reads |
|  | blaSHV | beta-lactamase | TP |  |  |  |
|  | blaOKP | beta-lactamase | TP | 50.00 | 100.00 |  |
| 90 mins |  |  |  |  |  | 2952 reads |
|  | blaLEN | beta-lactamase | TP | 66.67 | 100.00 |  |
| 120 mins |  |  |  |  |  | 3584 reads |
|  | oqxA | quinolone | TP | 83.33 | 100.00 |  |
| 570 mins |  |  |  |  |  | 8112 reads |
|  | oqxB | quinolone | TP | 100.00 | 100.00 |  |

### Comparison with other methods

To date, there are only a few existing pipelines for identification of species/subspecies from nanopore sequencing data, namely Metrichor [20], [8] and MetaPORE [9]. These methods commonly place the sample of question to a phylogeny taxonomy based on the number of reads that either are aligned to or have a similar k-mer profile to the taxon’s reference genome. Our species typing method is somewhat similar to this approach, although it additionally estimates confidence intervals in the species assignment. While we found that this approach can successfully identify species within 500 reads, the signal to noise from nanopore sequencing is too low to use a similar approach to correctly discriminate at the strain level, unless a large amount of data is available. Our strain typing uses a novel approach based on the presence and absence of genes and hence is able to make inference from a smaller number of reads.

Among the mentioned methods, only Metrichor [20] and MetaPORE [9] support genuine real-time analysis. As MetaPORE only focuses on viral species identification, we could only directly compare the performance of our method to Metrichor. We uploaded the first 1000 reads from our single samples and the first 3000 reads from our mixture sample to the Metrichor What’s In My Pot Bacteria k24 for SQK-MAP005 v1.27 (WIMP) workflow. Along with the species/subspecies and strains reported, WIMP provides a *classification score* filter where users can set the permissiveness of reporting. Table 5 presents the bacterial taxa reported by WIMP workflow for our data with the default classification score. For sample *K. pneumoniae* ATCC BAA-2146, WIMP only returned the taxon *K. pneumoniae* at the species level. On the other hand, for the second and third samples (*K. quasipneumoniae* ATCC 700603 and *K. pneumoniae* ATCC 13883), WIMP reported several *K. pneumoniae* strains but not the correct strain types of these samples (ST489 and ST3). For the mixture sample, two *E. coli* and three *S. aureus* strains were reported, but also not the correct strain types (*E. coli* ST73 and *S. aureus* ST243). While it was unclear whether the strain types of these samples were included in WIMP’s database, ST11 clearly was as it was reported in sample *K. pneumoniae* ATCC 700603. However, WIMP was unable to identify sample *K. pneumoniae* ATCC BAA-2146 to the strain level with 1000 reads, while our pipeline could do so in less than 400 reads (Figure 5).

**Table 5.** Report of Metrichor WIMP bacteria from the first 1000 reads of three single samples and the first 3000 reads of the mixture sample. The last column indicates if the detection is correct (✓) or incorrect (×) at species/strain levels. The Metrichor was able to identify the species (with some false positives) but not the strains in our samples.

| Sample | Reported by Metrichor | Strain type | Level | Accuracy |
| --- | --- | --- | --- | --- |
| *K. pneumoniae*<br>(ATCC BAA-2146, ST11) | *K. pneumoniae* | - | Species | ✓/ |
| *K. quasipneumoniae*<br>(ATCC 700603, ST489)* | *K. pneumoniae* subsp. *pneumoniae* | - | Sub-species | ✓/ |
|  | *K. pneumoniae* 342 | ST146 | Strain | ✓/× |
|  | *K. pneumoniae* JM45 | ST11 | Strain | ✓/× |
|  | *K. pneumoniae* CG43 | ST86 | Strain | ✓/× |
|  | *K. oxytoca* | - | Species | ×/ |
|  | *K. variicola* At-22 | - | Strain | ✓/× |
| *K. pneumoniae*<br>(ATCC 13883, ST3) | *K. pneumoniae* subsp. *pneumoniae* 1084 | ST1084 | Strain | ✓/× |
|  | *K. pneumoniae* CG43 | ST86 | Strain | ✓/× |
|  | *K. pneumoniae* subsp. *rhinoscleromatis* SB3432 | ST67 | Strain | ✓/× |
|  | *E. coli* O103:H2 str. 12009 | ST17 | Strain | ×/× |
| Mixture sample<br>75% *E. coli*<br>(ATCC 25922, ST73)<br>25% *S. aureus*<br>(ATCC 25923, ST243) | *E. coli* UMN026 | ST597 | Strain | ✓/× |
|  | *E. coli* ETEC H10407 | ST48 | Strain | ✓/× |
|  | *S. aureus* subsp. *aureus* HO 5096 0412 | ST22 | Strain | ✓/× |
|  | *S. aureus* subsp. *aureus* MRSA252 | ST36 | Strain | ✓/× |
|  | *S. aureus* subsp. *aureus* T0131 | ST239 | Strain | ✓/× |
|  | *Yersinia pestis* | - | Species | ×/ |

\**K. quasipneumoniae* ATCC 700603 strain was recently been re-classified from *K. pneumoniae* as *K. quasipneumoniae* but has not been updated in most major databases.

Antibiotic resistance genes detection from MinION sequencing was also explored in Judge et al [21]. Their approach was broadly similar to ours in that it initially aligns sequence reads to a resistance gene database, and then constructs a consensus sequence from the multiple alignment of matched reads. Both pipelines reported close to perfect resistance gene identification when compared with Illumina MiSeq sequencing. However, our pipeline uses a novel alignment parameter estimation using probabilistic Finite State Machines (see Methods). It is hence able to confidently report the presence of a resistance gene as soon as sufficient supporting data are available. This is the essence of real-time analysis presented here.

### Computational time

In our analyses, sequence reads were streamed through the pipeline in the exact order and timing as they were generated. Analysis results were generated periodically (every minute for species typing and strain typing and every five minutes for resistance gene identification). We examined the scalability of the pipeline to higher throughput by running the pipeline on a single computer equipped with 16 CPUs and streaming all sequence reads from the highest yield run (185Mb from sample *K. pneumoniae* ATCC BAA-2146) through the pipeline at 120 times higher speed than they were generated (*e.g.*, data sequenced in 2 minutes were streamed within 1 second). Analysis results were generated every 5 seconds for typing and every one minute for gene resistance analysis. With this hypothetical throughput, our pipeline correctly identified the species and strain of the sample in less than 20 seconds, upon which we could terminate the typing analyses. The pipeline then reported all the resistance genes in five minutes, which corresponded to the data generated in the first 10 hours of actual sequencing. This demonstrates the scalability of our pipeline to higher throughput sequencing platforms in the future.

### Real-time analysis of a clinical isolate

With the pipeline in place, we analysed a clinical *K. pneumoniae* isolate collected in Greece which was found to be resistant to a extensive range of antibiotics. We sequenced the sample on the MinlON with Chemistry R7.3 and ran the Metrichor service which performed basecalling and sample identification during the first three hours of the run. We also ran our pipeline in real-time on the base-called data returned from the Metrichor service.

We observed a delay from the base-calling of the data; the first read was sequenced on the MinlON within one minute from starting the run, but the basecalled data were received after 6 minutes. The delay tended to increase as more data were generated. We found the base-called data returned during the three hour run of the Metrichor service were actually sequenced within 45 minutes on the MinION. This highlights the need for a local base-calling step to improve real-time analysis. Figures 6a and 6b show the timing (from the start of the MinlON run) of sample identification using our pipeline. The pipeline reported *K. pneumoniae* as the only species in the sample within 10 minutes, and reached a confidence interval of less than 0.1 in 40 minutes when approximately 200 reads were analysed. We noticed that these 200 reads were actually sequenced in 7 minutes by the MinION. For strain identification, our pipeline initially reported ST1199 but after 2.5 hours, reported ST258 as the sequence type for this isolate. It is worth noting that the two strains are highly similar; their MLST profiles differ by only one SNP in the seven housekeeping genes. By sequencing the isolate on the Illumina MiSeq as described above, we confirmed that the sequence type for the strain is ST258. On the other hand, the sample identification from Metrichor initially reported *K. pneumoniae* 1084 (ST23), but finally reported two strains namely *K. pneumoniae* JM45 (ST11) and *K. pneumoniae* HS11286 (ST11) after 3 hours (Figure 6c). During the three hour run with less than 4000 reads (16Mb of data), our pipeline reported two antibiotic resistance genes, namely sul2 (sulphonamide) and tetA (tetracycline). Our analysis of the Illumina data for this strain confirmed the presence of these two genes. Clinical susceptibility testing also showed the resistance of this isolate to tetracycline and Sulfamethoxazole-trimethoprim (MIC ≥ 16 *μ*g/mL and ≥ 320 *μ*g/mL, respectively analyzed by VITEK©2 bioMerieux, Inc). Finally, we re-analysed the data from this run using the emulation described previously, and obtained the same results as from the real-time analysis.

**Figure 6.**
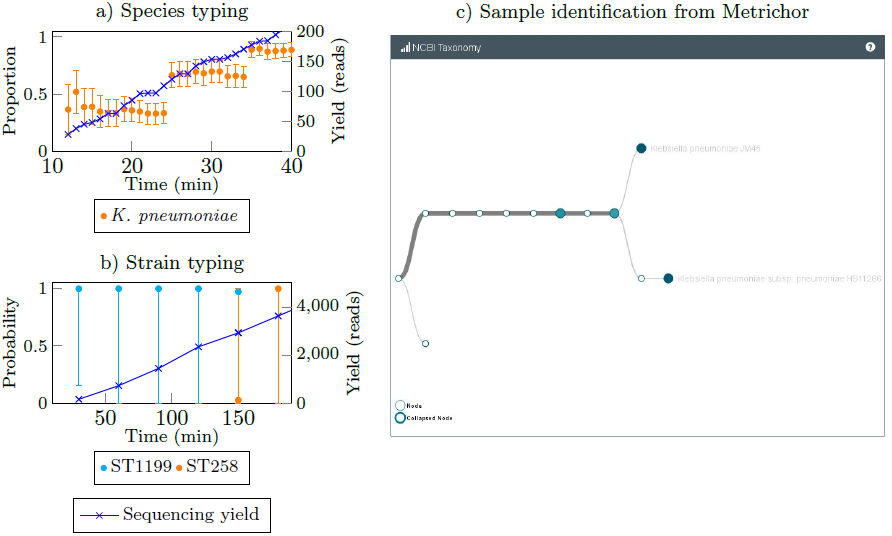
Real-time sample identification of a clinical isolate directly from the MinION using our pipeline and by Metrichor service. The time includes basecalling timing.

## Discussion

In recent years HTS has become an integrative tool for infectious disease research [22, 23]. There have been several reports emphasizing the use of HTS methods to characterize clinical isolates, to study the spread of drug resistant microorganisms and to investigate outbreak of infections [24–26]. These studies predominantly use massively parallel short-read sequencing technologies such as the Illumina Miseq, NextSeq or HiSeq. These sequencers achieve a very high base calling accuracy which makes them ideally suited to applications which require accurate calling of single nucleotide polymorphisms (SNPs), including reconstructing the evolutionary history of different bacterial isolates; tracking transmissions during an outbreak; placing a new isolate on a phylogenetic tree and population genetic analyses. However, these technologies attain their high yield by sequencing a single base per cycle for millions of sequence fragments in parallel, where each cycle takes at least 5 minutes.

The Oxford Nanopore MinION device, on the other hand, generated as many as 500 reads in the first 10 minutes of sequencing in our hands (which is 3 times lower than the theoretical maximum). The error rate of these reads was substantially higher than the corresponding Illumina data. Existing bioinformatics algorithms - which have been developed for highly accurate Sanger and subsequently for short-read sequencing - rely on accurate base and SNP calling, which makes their application to MinION data challenging. As an example, most existing strain typing approaches often use a MLST system, either on a predefined set of house keeping genes [27], or on core genes set [28]. These approaches are highly standardized, reproducible and portable, and hence are routinely used in laboratories around the world. Rapid genomics diagnosis tools using MLST from high-throughput sequencing such as SRST2 [29] have also been developed. While we showed that MLST can be adapted to identify bacterial strain type from nanopore sequencing, this requires high coverage sequencing of the gene set to overcome the high error rates. Similarly, other researchers have shown that error correction can overcome the high error rate providing enough coverage is obtained [15, 30].

The main contribution of this manuscript is to demonstrate that despite the higher error rate, it is possible to return clinical actionable information, including species and strain types from as few as 500 reads. We achieved this by developing novel approaches which are less sensitive to base-calling errors and which use whatever subset of genome-wide information is observed up to a point in time, rather than a panel of pre-defined markers or genes. For example, the strain typing presence/absence approach relies only on being able to identify homology to genes and also allows for a level of incorrect gene annotation.

Our species typing module has some similarities to the approach used by MetaPhlAn [31], in that we use the proportion of reads which map to different taxonomic groupings to estimate the proportion of different species in a sample. MetaPhlan optimises computational speed by aligning to a precomputed database of sequences which are pervasive within a single taxonomic grouping but not seen outside that grouping. This allows it to blast against a database which is 20 times smaller than a full bacterial genomic database. This was designed to make metagenomics inference feasible on datasets with millions of reads. On the other hand, our species typing approach is designed to make a similar inference using only hundreds of reads, and moreover, also continuously updates confidence intervals so the user knows when they can stop sequencing and make a diagnosis.

Our strain typing module has the advantage of being able to rapidly type a known strain with a small number of low quality (*i.e.* mostly 1D) reads. Competiting approaches which use kmers, such as that implemented in ‘WIMP’ appear to require substantially more high quality data. The drawback of our approach is that if a large number of genes are lost or gained in a single event, such as the gain or loss of a plasmid, the most likely strain may be incorrect, although the bootstrap-derived confidence intervals will be wide in this case.

Our antibiotic resistance module is able to identify the drug resistance potential of an isolate within a few hours of sequencing with very high specificity. In particular, with the most recent chemistry utilized in this paper (R7.3), we were able to identify the complete resistance potential of a *K. pneumoniae* isolate without any false positives in 9.5 hours and with approximately 8000 reads (80% of the resistance genes were identified with 3000 reads in 2 hours). In order to achieve high specificity we designed a probabilistic Finite State Machine for error correction. This approach continuously updates the consensus sequence from the multiple alignment of reads, and re-estimates the error profile of the consensus sequence. This allows the reporting of the presence of a resistance gene once sufficient accuracy is obtained, rather than waiting for the full run to complete.

In summary, we have developed an open-source, flexible pipeline for real-time analysis of MinION sequencing data. The only step in our pipeline in which data are written to, and then re-read from disk is the base-calling step using Metrichor. npReader immediately identifies new reads as they are generated by Metrichor, however some delay can occur due to waiting for base-called data to be returned from Metrichor. Oxford Nanopore Technologies have recently opened up the Application Programming Interface to extract raw data directly from the MinION. This, together with the recent development of open source base-calling algorithms [32, 33] to run on the local machine, will allow future development of a completely streaming pipeline, in the sense of never saving data to disk. Our pipeline can be deployed on a single 16 core computer, capable of analysing MinION data streaming at up to 120x the current rate of sequencing; or on a high performance computing cluster to scale with the potential even higher throughput of forthcoming nanopore sequencing platforms. Our pipeline incorporates three streaming algorithms, but further algorithms can be flexibly integrated into this pipeline. Other algorithms such as for base-calling [32, 33] or alignment of signal-level data can also be integrated into the pipeline to by-bass the base-calling currently from Metrichor.

Other investigators have focused on the long-read nature of MinION sequencing data, which enables complete genome assembly [30] as well as the identification of sites of integration of resistance islands [13]. Researchers have also recently reported that MinION sequencing data could accurately identify bacterial outbreak strains within 50 minutes of sequencing [8] by placing reads onto a phylogenetic tree; and drug resistance profile of a *S. aureus* sample determined using a de-Bruijn graph approach from 8 hours of sequencing data [34].

We have shown that switching from a traditional short-read sequencing pipeline coupled with standard, non-streaming bioinformatics algorithms, to a nanopore sequencing pipeline coupled with streaming bioinformatics algorithms can dramatically cut the time taken from DNA library to results from at least 8 hours down to 30 minutes. With the time for library preparation for nanopore sequencing forecast to be shortened to 10 minutes, the major time bottleneck then becomes the bacterial culture step (which can be 24 hours). The MinION sequencer can be used on clinical sample without culture, however this then dilutes the proportion of bacterial DNA present. Nevertheless, this may become a viable time-sensitive strategy as sequencing yield increases, particularly with high colony-forming-unit (CFU) infections. Another promising approach may be to use approaches to pre-concentrate bacterial DNA [35].

One of the major advantages of a whole-genome sequencing approach to drug resistance profiling is that it is not necessary to restrict the analysis to a limited panel of drug-resistance tests but it is possible to discover the complete drug resistance profile in a sample. With a complete picture of the drug-resistance profile within a few hours, a clinician may be able to design an antibiotic treatment regimen that is both more likely to succeed and less likely to induce further antibiotic resistance. However, even achieving completely accurate identification of resistance genes is only a first step in accurately predicting the resistance profile, as mutations may effect the rate at which these genes are transcribed and also their antibiotic resistance activity. Prediction of antibiotic resistance from genotype is an area which warrants substantial further research.

## Methods

### DNA extraction

Bacterial strains *K. pneumoniae* ATCC BAA-2146, ATCC 13883, *K. quasipneumoniae* ATCC 700603, *E. coli* ATCC 25922 and *S. aureus* ATCC 25923 were obtained from American Type Culture Collection (ATCC, USA). *K. pneumoniae* clinical isolate was acquired from Hygeia General Hospital, Athens, Greece from a patient stool sample in 2014 (Lab ID 100575214, isolate 1). Clinical susceptibility profiling by VITEK®2 (bioMérieux Inc.) identified the isolate as Carbapenemase producing (KPC) giving rise to extended spectrum *β*-lactam resistance. It was also deemed resistant to aminoglycoside, phenicol, quinolone, sulphonamide, tetracycline and trimethoprim antibiotics, rendering it an extensively drugresistant bacterial isolate. Bacterial cultures were grown overnight from a single colony at 37° C with shaking (180 rpm). Whole cell DNA was extracted from the cultures using the DNeasy Blood and Tissue Kit (QIAGEN©, Cat #69504) according to the bacterial DNA extraction protocol with enzymatic lysis pre-treatment.

### MinION sequencing

Library preparation was performed using the Genomic DNA Sequencing kit (Oxford Nanopore) according to the manufacturer’s instruction. For the R7 MinION Flow Cells SQK-MAP-002 sequencing kit was used and for R7.3 MinION Flow Cells SQK-MAP-003 or SQK-MAP-006 Genomic Sequencing kits were used according to the manufacturer’s instruction.

For the library mixture sample, DNA concentration of each library was measured using Qubit Fluorimeter (Thermo Fisher Scientific). Based on the concentration, 75% of *E. coli* (ATCC 25922) library and 25% of *S. aureus* (ATCC 25923) library were mixed prior to sequencing.

For each sample a new MinION Flow Cell (R7 or R7.3) was used for sequencing. The library mix was loaded onto the MinION Flow Cell and the Genomic DNA 48 hour sequencing protocol was initiated on the MinKNOW software.

### MinION data analysis

The sequence read data were base called with Metrichor Agent (

~~~
https://metrichor.com
~~~

). We used npReader [10] to convert base-called sequence data in fast5 format to fastq format. The npReader program also extracted the time that each read was sequenced and used this information to sort the read sequences in order they were produced. For the real-time analyses, we wrote a program to emulate the sequencing process in that it streamlined each read in the exact order it was sequenced. The program also allowed scaling up the sequencing emulation to a factor of choice. Our pipeline allows for filtering out 1D reads at multiple stages (including via npReader). All subsequent analyses in this paper used both 1D and 2D reads.

### MiSeq sequencing and data analysis

Library preparation was performed using the NexteraXT DNA Sample preparation kit (Illumina) as recommended by the manufacturer. Libraries were sequenced on the MiSeq instrument (Illumina) with 300bp paired end sequencing, to a coverage of over 100-fold. Read data were trimmed with *trimmomatic* [36] (V0.32) and subsequently assembled using SPAdes [37] (V3.5), resulting in assemblies with N50 exceeding 200kb. Their strain types were identified by submitting the assembled genomes to the MLST servers 

~~~
https://cge.cbs.dtu.dk/services/MLST/
~~~

 [38] for *K. pneumoniae, E. coli* (set #1) and *S. aureus*.

We identified the antibiotic resistance profiles of these strains from their MiSeq assemblies. We used blastn (V2.29) to align these assemblies to the database of resistance genes obtained from ResFinder [17]. Genes which were covered at least greater than 85% by the alignments and with greater than 85% sequence identity were considered to be present in the sample. These gene profiles were used as a benchmark to validate the MinION sequencing analysis.

### Species typing

We downloaded the bacterial genome database on GenBank (

~~~
ftp://ftp.ncbi.nlm.nih.gov/genomes/Bacteria/
~~~

, accessed 19 Nov 2014), which contained high quality complete genomes of 2785 bacterial strains from 1487 bacteria species. We expanded this database to include two *K. quasipneumoniae* genomes. Our species typing pipeline streamed read data from npReader directly to BWA-MEM [11] (V0.7.10-r858) which aligned the reads to the database. Output from BWA in SAM format was streamed directly into our species typing pipeline, which calculated the proportion of reads aligned to each of these species. Our species typing method considers the proportions {*p*_1_,*p*_2_,…, *P_k_* } of k species in the mixture as the parameters of a k-category multinomial distribution, and the read counts {*c*_1_, *c*_2_,…,*c_k_* } for the species as an observation from *c*_1_ + *c*_2_ + … + *c_k_* independent trials drawn from the distribution. It then uses the MultinomialCI package in R [39] to calculate the 95% confidence intervals of these proportions from the observation.

### MLST typing

MinION sequence reads from *K. pneumoniae* strains were aligned to the seven house-keeping genes specified by the MLST system using BWA-MEM [11]. We then collected reads that were aligned to a gene and performed a multiple alignment on them using kalign2 [40]. The consensus sequence created from the multiple alignment was then globally aligned to all alleles of the gene using a probabilistic Finite State Machine (see below) for global alignment. The score of a MLST type was determined by the sum of the scores of seven alleles making up the type.

### Strain typing

We built gene profile databases for *K. pneumoniae, E. coli* and *S. aureus* from the RefSeq annotation. Specifically, we obtained the publicly available assemblies of these species listed on RefSeq master database (

~~~
ftp://ftp.ncbi.nih.gov/genomes/ASSEMBLY_REPORTS/assembly_summary_refseq.txt
~~~

, accessed 17 July 2015). We used the relevant MLST schemes obtained from (

~~~
https://cge.cbs.dtu.dk/services/MLST/
~~~

 [38] to identify strain type of each assembly. For each strain type, we selected the assembly with highest N50 statistic and use the RefSeq gene annotation of the assembly to determine the gene content of the strain type.

In order to develop a simple probabilistic presence/absence strain typing model, we consider the genomes of each of the strains simply as a collection of genes. Denote by *St*_*j*=1…*J*_ the all the strains in our database (for a fixed species). Denote by *g*_*j*,*k*_ the *k*^*th*^ gene in the database for strain j, where the genes are listed in no particular order. Denote by *N_j_* the total number of genes in *St_j_*.

We align each sequence read *r_i_* from the MinION device to the gene database using BWA-MEM [11]. We count the number of genes of each strain that are aligned to some reads, denoted *N_j_*(*r_i_*).

We describe below how we can calculate a likelihood, *P*(*r_i_* |*St_j_*), of each strain generating each read, from which we can calculate the posterior probability of each strain *St_j_* conditional on observing the reads *r*_1_…*r_m_*:

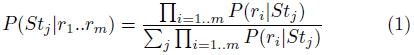

The probability P(*r_i_* |*St_j_*) could be calculated using a simple model as

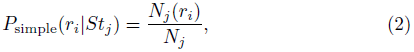

however, this model suffers from the problem that if we observe any read which overlaps a gene not in the reference genome for Stj, then the posterior probability of that strain will become zero. Thus, this model is very unstable. In order to make this estimate more stable, we use a mixture model which allows for the read to have been generated by a background model:

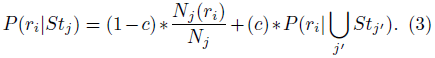

The background model considers the probability that the read was generated from any of the strains,

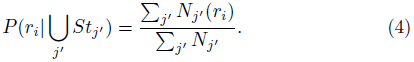

This makes the posterior probability estimates more stable. It also makes the model robust to incorrect annotation of the reads from the MinION sequencer and incorrect annotation of the reference genome. We have investigated use of *c* = 0.2, *c* = 0.1 and *c* = 0.05 and found that it has little impact on the results, with slightly smaller confidence intervals (data not shown). We choose *c* = 0.2 in order to conservatively estimate confidence intervals.

Finally, in order to calculate confidence intervals we employ a bootstrap resampling approach in which we resample *m* reads from *r*_1_,…*r_m_* with replacement. This is repeated 1000 times, and the posterior probabilities are recalculated every iteration. We calculate the 95% confidence intervals from the empirical distribution of these posterior probabilities.

To gain some insight into how this model works in response to gene presence, consider a gene *g* which is present in a fraction *f* of strains, including *St_j_* but not including *St*_k_. For simplicity assume that each strain has *N* genes. The difference in log-likelihood *St_j_* and *St_k_* conditional on *g* can be approximated by log(1/*c*) + log(1/*f*), showing that a more specific gene has a stronger effect in our model than a common gene in distinguishing strains.

To gain insight into the effect of gene absence in contrast to gene presence, assume instead that the only difference between *St_j_* and St_k_ is that a single gene (*g*) is deleted in *St_j_*, and denote by *N* = *N_j_* = *N_k_* – 1. If we sequence *N* ln(2) genes from *St_j_* without seeing gene g, the difference in log-likelihood becomes *N* ln(2) * (log(*N*) – log(*N* – 1)) ≈ 1 bit, corresponding to the likelihood for *St_j_* being twice as big as the likelihood of *St_k_*. For example, if a strain has 1000 genes, then we would need to observe 693 genes without observing g to be able to conclude that the observed data were twice as likely to be generated from the species with a single gene deletion. For comparison, we would need to only sequence 100 genes from *St_k_* to get an expected log-likelihood difference of 1 bits versus *St_j_*, demonstrating the extra information in gene ‘presence’ versus ‘absence’ typing.

### Antibiotic resistance gene classes detection

We downloaded the resistance gene database from Res-Finder(

~~~
https://cge.cbs.dtu.dk/services/ResFind
~~~

 accessed July 2015). We aligned each gene to the collection of bacterial genomes in RefSeq using blastn [41], and used the best alignment of the gene to extract 100 bp sequences flanking the antibiotic resistance genes. We found that the inclusion of these flanking sequences improved the sensitivity of mapping MinION reads to the gene database.

We then grouped these genes based on 90% sequence identity into 609 groups. We manually checked and found genes within a group were variants of the same gene. We selected the longest gene in each group to make up a reduced resistance gene database. To create a benchmark of resistance gene for a sample, we blastn the Illumina assembly of the sample against this reduced gene database, and reported genes with greater than 85% coverage and identity.

Our analysis pipeline aligned MinION sequencing data into this reduced resistance gene database using BWA-MEM [11] in a streamline fashion, and examined genes that had reads mapping to the whole gene (not including flanking sequences). Due to the high error rates of MinION sequence data, we noticed a high rate of false positive genes. To reduce false positives, we used kalign2 [40] to perform a multiple alignment of reads that were aligned to the same gene. The consensus sequence resulting from the multiple alignment was then compared with the gene sequence using a probabilistic Finite State Machine (see below). The pipeline then reported gene classes based on the genes detected.

### Sensitive alignment of noisy sequences with probabilistic Finite State Machines

Our methods for MLST strain typing and antibiotic resistance gene identification require the alignment of a consensus sequence to a gene or a gene allele. Such an alignment generally assumes a model and a set of parameters of the differences between the sequences. It is widely recognised that the accuracy of the alignment is sensitive to these parameters [42–44]. However, in the context of real-time analysis of MinION sequencing, it is not possible to select in advance a sensible set of parameters. On the one hand, the quality among sequence reads differs remarkably; as shown in Figure 3 and Table 2 – the majority (95%) of the reads across our four runs have the Phred score ranging between 3 and 7 for template and complement reads (corresponding to 50% - 80% accuracy) and between 6 to 12 for 2D reads (75%-95% accuracy). On the other hand, a consensus sequence is computationally constructed from a set of reads. Its quality is hence contingent to not only the quality of the reads but also the number of reads in the set.

We use a probabilistic Finite State Machine (pFSM) [45 rt/o, model the differences, and hence the simultaneous error profile of the consensus sequence. Briefly, a pFSM is a probabilistic model of genomic alignment that takes into acount different types of variations including SNPs, insertions and deletions. A pFSM is equivalent to a hidden Markov Model. The pFSM consists of a set of states and transitions between states. Each transition corresponds to an *action* and is associated with a cost for the action. An action could be one of *copy* (*C*), *substitute* (*S*), *delete* (*D*) and *insert* (*I*). Figure 7 depicts a three-state pFSM which is equivalent to a affine gap penalty alignment model. In order to assess an alignment of two sequences A and B, under a hypothesis specified by the parameters, the pFSM computes the cost to generate one sequence (say A) given the other (B). For example, while in state *Copy*, the machine consumes the next base in B, generates the next base in A; it is said to take action *C* if the two bases are the same, or action *S* otherwise, and to follow either transition to state *Copy*. Alternatively, the machine can take either action *D* (consumes the next base in B without generating any base in A and moves to state *Delete)*, or action *I* (generates the next base in A without consuming a base in B and moves to state *Insert*). These actions are repeated until the whole sequence B is generated.

We use an information-theoretic measure where the cost of a transition is that of encoding the generated base, or in other words, the negative logarithm of the probability of the associated action (*c* = –*log*_2_(*P*(*a*)). The foundation of this approach goes back to the 1960s when it was proposed as a basis for inductive inference [46, 47]. It has since been used in a number of bioinformatics applications such as for calculating the BLOSUM matrix [48] and modelling DNA sequences [49, 50]. More importantly, this information-theoretic framework allows one to estimate a sensible set of parameters for any *related* two sequences. This is done via a Expectation-Maximisation process. This starts with an initial set of probabilities at each state. In the E-step, the best alignment (lowest cost) is calculated by a dynamic programming algorithm. The frequencies of actions at each state are then used to re-estimate the probabilities in the M-step. A detailed discussion of this process is provided in Allison et al [45] and Cao et al [51]. The process is guaranteed to converse to an optimal, and it does so in only a few iterations in our experience.

**Figure 7.**
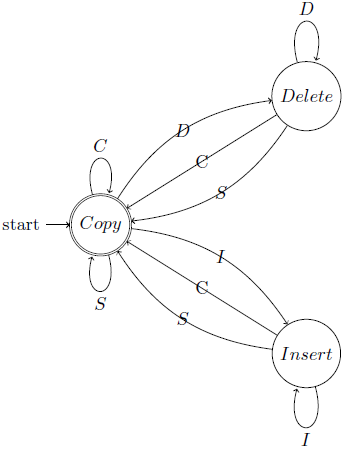
Schematic of a three-state probabilistic Finite State Machine.

## Availability of supporting data

All scripts for the presented analyses are provided in 

~~~
https://github.com/mdcao/npAnalysis
~~~

. The source code of the software is publicly available in github repository (

~~~
https://github.com/mdcao/japsa
~~~

). The MinION sequencing data for the three single samples are available in European Nucleotide Archive Study Accession Number ERP010377 (

~~~
http://www.ebi.ac.uk/ena/data/view/ERP010377
~~~

). The MinION sequencing data for mixture sample and the Illumina sequencing are in the process of depositing to European Nucleotide Archive. They are made available via the links provided in 

~~~
https://github.com/mdcao/npAnalysis
~~~

.

## List of abbreviations used

MLST: Multilocus sequence typing
pFSM: probabilistic Finite State Machine.

## Author’s contributions

MDC, DG, MC and LC conceived the study, performed the analysis and wrote the first draft of the manuscript. AE performed the bacterial cultures and DNA extractions. DG performed the MinION sequencing. MDC and LC designed and developed the algorithms and the analysis framework. MDC, HZ, and LC performed the bioinformatics analyses. All authors contributed to editing the final manuscript.

## Acknowledgements

We thank Ilias Karaiskos and Helen Giamarellou (6th Dept. of Internal Medicine, Hygeia General Hospital, Athens, Greece) for providing the clinical *K. pneumoniae* isolate. MAC is an NHMRC Principal Research Fellow (APP1059354). LC is an ARC Future Fellow (FT110100972). The research is supported by funding from the Institute for Molecular Bioscience Centre for Superbug Solutions (610246).

## References

1. Boyd, S.D.: Diagnostic applications of high-throughput DNA sequencing. Annual review of pathology 8, 381–410 (2013). doi:10.1146/annurev-pathol-020712-164026

2. Koboldt, D.C., Steinberg, K.M., Larson, D.E., Wilson, R.K., Mardis, E.R.: The next-generation sequencing revolution and its impact on genomics. Cell 155(1), 27–38 (2013). doi:10.1016/j.cell.2013.09.006

3. Gaber, M.M., Zaslavsky, A., Krishnaswamy, S.: Mining data streams. ACM SIGMOD Record 34(2), 18 (2005). doi:10.1145/1083784.1083789

4. Muthukrishnan, S.: Data Streams: Algorithms and Applications. Foundations and Trends in Theoretical Computer Science 1(2), 117–236 (2005)

5. Kasianowicz, J.J., Brandin, E., Branton, D., Deamer, D.W.: Characterization of individual polynucleotide molecules using a membrane channel. Proceedings of the National Academy of Sciences 93(24), 13770–13773 (1996). doi:10.1073/pnas.93.24.13770

6. Branton, D., Deamer, D.W., Marziali, A., Bayley, H., Benner, S.A., Butler, T., Di Ventra, M., Garaj, S., Hibbs, A., Huang, X., Jovanovich, S.B., Krstic, P.S., Lindsay, S., Ling, X.S., Mastrangelo, C.H., Meller, A., Oliver, J.S., Pershin, Y.V., Ramsey, J.M., Riehn, R., Soni, G.V., Tabard-Cossa, V.., Wanunu, M., Wiggin, M., Schloss, J.A.: The potential and challenges of nanopore sequencing. Nature biotechnology 26(10), 1146–53 (2008). doi:10.1038/nbt.1495

7. Stoddart, D., Heron, A.J., Mikhailova, E., Maglia, G., Bayley, H.: Single-nucleotide discrimination in immobilized DNA oligonucleotides with a biological nanopore. Proceedings of the National Academy of Sciences of the United States of America 106(19), 7702–7 (2009). doi:10.1073/pnas.0901054106

8. Quick, J., Ashton, P., Calus, S., Chatt, C., Gossain, S., Hawker, J., Nair, S., Neal, K., Nye, K., Peters, T., De Pinna, E., Robinson, E., Struthers, K., Webber, M., Catto, A., Dallman, T.J., Hawkey, P., Loman, N.J.: Rapid draft sequencing and real-time nanopore sequencing in a hospital outbreak of Salmonella. Genome Biology 16(1), 114 (2015). doi:10.1186/s13059-015-0677-2

9. Greninger, A.L., Naccache, S.N., Federman, S., Yu, G., Mbala, P., Bres, V., Stryke, D., Bouquet, J., Somasekar, S., Linnen, J.M., Dodd, R., Mulembakani, P., Schneider, B.S., Muyembe-Tamfum, J.-J., Stramer, S.L., Chiu, C.Y.: Rapid metagenomic identification of viral pathogens in clinical samples by real-time nanopore sequencing analysis. Genome Medicine 7(1), 99 (2015). doi:10.1186/s13073-015-0220-9

10. Cao, M.D., Ganesamoorthy, D., Cooper, M.A., Coin, L.J.M.: Realtime analysis and visualization of MinION sequencing data with npReader. Bioinformatics 32(5), 764–766 (2016). doi:10.1093/bioinformatics/btv658

11. Li, H.: Aligning sequence reads, clone sequences and assembly contigs with BWA-MEM, 3 (2013). 1303.3997#

12. Quick, J., Quinlan, A.R., Loman, N.J.: A Reference Bacterial Genome Dataset Generated on the {MinION } Portable Single-molecule Nanopore Sequencer. GigaScience 3(1), 22 (2014). doi:10.1186/2047-217x-3-22

13. Ashton, P.M., Nair, S., Dallman, T., Rubino, S., Rabsch, W., Mwaigwisya, S., Wain, J., O’Grady, J.: MinION nanopore sequencing identifies the position and structure of a bacterial antibiotic resistance island. Nature Biotechnology 33(3), 296–300 (2015). doi:10.1038/nbt.3103

14. Kilianski, A., Haas, J.L., Corriveau, E.J., Liem, A.T., Willis, K.L., Kadavy, D.R., Rosenzweig, C.N., Minot, S.S.: Bacterial and viral identification and differentiation by amplicon sequencing on the MinION nanopore sequencer. GigaScience 4(1) (2015). doi:10.1186/s13742-015-0051-z

15. Jain, M., Fiddes, I.T., Miga, K.H., Olsen, H.E., Paten, B., Akeson, M.: Improved data analysis for the MinION nanopore sequencer. Nature Methods 12(4), 351–356 (2015). doi:10.1038/nmeth.3290

16. Diancourt, L., Passet, V., Verhoef, J., Grimont, P.A.D., Brisse, S.: Multilocus Sequence Typing of Klebsiella pneumoniae Nosocomial Isolates. Journal of Clinical Microbiology 43(8), 4178–4182 (2005). doi:10.1128/JCM.43.8.4178-4182.2005

17. Zankari, E., Hasman, H., Cosentino, S., Vestergaard, M., Rasmussen, S., Lund, O., Aarestrup, F.M., Larsen, M.V.: Identification of Acquired Antimicrobial Resistance Genes. Journal of Antimicrobial Chemotherapy 67(11), 2640–2644 (2012). doi:10.1093/jac/dks261

18. Allison, L., Wallace, C.S., Yee, C.N.: When is a String Like a String? In: International Symposium on Artificial Intelligence and Mathematics (1990)

19. Poznik, D.G., Henn, B.M., Yee, M.-C., Sliwerska, E., Euskirchen, G.M., Lin, A.A., Snyder, M., Quintana-Murci, L.., Kidd, J.M., Underhill, P.A., Bustamante, C.D.: Sequencing {Y } Chromosomes Resolves Discrepancy in Time to Common Ancestor of Males Versus Females. Science 341(6145), 562–565 (2013). doi:10.1126/science.1237619

20. Juul, S., Izquierdo, F., Hurst, A., Dai, X., Wright, A., Kulesha, E., Pettett, R., Turner, D.J.: What’s in my pot? Real-time species identification on the MinION. bioRxiv (2015). doi:10.1101/030742

21. Judge, K., Harris, S.R., Reuter, S., Parkhill, J., Peacock, S.J.: Early insights into the potential of the Oxford Nanopore MinION for the detection of antimicrobial resistance genes. Journal of Antimicrobial Chemotherapy 70(10), 2775–2778 (2015). doi:10.1093/jac/dkv206

22. Dunne, W.M., Westblade, L.F., Ford, B.: Next-generation and whole-genome sequencing in the diagnostic clinical microbiology laboratory. European journal of clinical microbiology & infectious diseases: official publication of the European Society of Clinical Microbiology 31(8), 1719–26 (2012). doi:10.1007/s10096-012-1641-7

23. Fricke, W.F., Rasko, D.A.: Bacterial genome sequencing in the clinic: bioinformatic challenges and solutions. Nature reviews. Genetics 15(1), 49–55 (2014). doi:10.1038/nrg3624

24. Hudson, C.M., Bent, Z.W., Meagher, R.J., Williams, K.P.: Resistance determinants and mobile genetic elements of an NDM-1-encoding Klebsiella pneumoniae strain. PloS one 9(6), 99209 (2014). doi:10.1371/journal.pone.0099209

25. Petty, N.K., Ben Zakour, N.L., Stanton-Cook, M.., Skippington, E., Totsika, M., Forde, B.M., Phan, M.-D., Gomes Moriel, D., Peters, K.M., Davies, M., Rogers, B.A., Dougan, G., Rodriguez-Bano, J.., Pascual, A., Pitout, J.D.D., Upton, M., Paterson, D.L., Walsh, T.R., Schembri, M.A., Beatson, S.A.: Global dissemination of a multidrug resistant Escherichia coli clone. Proceedings of the National Academy of Sciences 111(15), 5694–5699 (2014). doi:10.1073/pnas.1322678111

26. Stoesser, N., Giess, A., Batty, E.M., Sheppard, a.E., Walker, a.S., Wilson, D.J., Didelot, X., Bashir, A., Sebra, R., Kasarskis, A., Sthapit, B., Shakya, M., Kelly, D., Pollard, a.J., Peto, T.E.a., Crook, D.W., Donnelly, P., Thorson, S., Amatya, P., Joshi, S.: Genome Sequencing of an Extended Series of NDM-Klebsiella pneumoniae Neonatal Infections in a Nepali Hospital Characterizes the Extent of Community Versus Hospital-associated Transmission in an Endemic Setting. Antimicrobial agents and chemotherapy 58(12), 7347–57 (2014). doi:10.1128/AAC.03900-14

27. Maiden, M.C., Bygraves, J.a., Feil, E., Morelli, G., Russell, J.E., Urwin, R., Zhang, Q., Zhou, J., Zurth, K., Caugant, D.a., Feavers, I.M., Achtman, M., Spratt, B.G.: Multilocus sequence typing: a portable approach to the identification of clones within populations of pathogenic microorganisms. Proceedings of the National Academy of Sciences of the United States of America 95(6), 3140–3145 (1998). doi:10.1073/pnas.95.6.3140

28. Cody, A.J., McCarthy, N.D., Jansen van Rensburg, M., Isinkaye, T., Bentley, S.D., Parkhill, J., Dingle, K.E., Bowler, I.C.J.W., Jolley, K.A., Maiden, M.C.J.: Real-Time Genomic Epidemiological Evaluation of Human Campylobacter Isolates by Use of Whole-Genome Multilocus Sequence Typing. Journal of Clinical Microbiology 51(8), 2526–2534 (2013). doi:10.1128/JCM.00066-13

29. Inouye, M., Dashnow, H., Raven, L.-A., Schultz, M.B., Pope, B.J., Tomita, T., Zobel, J., Holt, K.E.: SRST2: Rapid genomic surveillance for public health and hospital microbiology labs. Genome medicine 6(11), 90 (2014). doi:10.1186/s13073-014-0090-6

30. Loman, N.J., Quick, J., Simpson, J.T.: A complete bacterial genome assembled de novo using only nanopore sequencing data. Nature Methods 12(8), 733–735 (2015). doi:10.1038/nmeth.3444

31. Segata, N., Waldron, L., Ballarini, A., Narasimhan, V., Jousson, O., Huttenhower, C.: Metagenomic microbial community profiling using unique clade-specific marker genes. Nature methods 9(8), 811–4 (2012). doi:10.1038/nmeth.2066

32. David, M., Dursi, L.J., Yao, D., Boutros, P.C., Simpson, J.T.: Nanocall: An Open Source Basecaller for Oxford Nanopore Sequencing Data. bioRxiv, 046086 (2016). doi:10.1101/046086

33. BoZa, V., Brejova, B., Vinar, T.: DeepNano: Deep Recurrent Neural Networks for Base Calling in MinION Nanopore Reads (2016). 1603. 09195

34. Bradley, P., Gordon, N.C., Walker, T.M., Dunn, L., Heys, S., Huang, B., Earle, S., Pankhurst, L.J., Anson, L., de Cesare, M., Piazza, P., Votintseva, A.A., Golubchik, T., Wilson, D.J., Wyllie, D.H., Diel, R., Niemann, S., Feuerriegel, S., Kohl, T.A., Ismail, N., Omar, S.V., Smith, E.G., Buck, D., McVean, G., Walker, A.S., Peto, T.E.A., Crook, D.W., Iqbal, Z.: Rapid antibiotic-resistance predictions from genome sequence data for Staphylococcus aureus and Mycobacterium tuberculosis. Nature Communications 6, 10063 (2015). doi:10.1038/ncomms10063

35. Hasman, H., Saputra, D., Sicheritz-Ponten, T.., Lund, O., Svendsen, C. A., Frimodt-Møller, N., Aarestrup, F.M.: Rapid whole-genome sequencing for detection and characterization of microorganisms directly from clinical samples. Journal of clinical microbiology 52(1), 139–46 (2014). doi:10.1128/JCM.02452-13

36. Bolger, A.M., Lohse, M., Usadel, B.: Trimmomatic: a flexible trimmer for Illumina sequence data. Bioinformatics 30(15), 2114–2120 (2014). doi:10.1093/bioinformatics/btu170

37. Bankevich, A., Nurk, S., Antipov, D., Gurevich, A.A., Dvorkin, M., Kulikov, A.S., Lesin, V.M., Nikolenko, S.I., Pham, S., Prjibelski, A.D., Pyshkin, A.V., Sirotkin, A.V., Vyahhi, N., Tesler, G., Alekseyev, M.A., Pevzner, P.A.: SPAdes: A New Genome Assembly Algorithm and Its Applications to Single-Cell Sequencing. Journal of Computational Biology 19(5), 455–477 (2012). doi:10.1089/cmb.2012.0021

38. Larsen, M.V., Cosentino, S., Rasmussen, S., Friis, C., Hasman, H., Marvig, R.L., Jelsbak, L., Sicheritz-Ponten, T.., Ussery, D.W., Aarestrup, F.M., Lund, O.: Multilocus Sequence Typing of Total-Genome-Sequenced Bacteria. Journal of Clinical Microbiology 50(4), 1355–1361 (2012). doi:10.1128/JCM.06094-11

39. Sison, C.P., Glaz, J.: Simultaneous Confidence Intervals and Sample Size Determination for Multinomial Proportions. Journal of the American Statistical Association 90(429), 366 (1995). doi:10.2307/2291162

40. Lassmann, T., Frings, O., Sonnhammer, E.L.L.: Kalign2: High-performance Multiple Alignment of Protein and Nucleotide Sequences Allowing External Features. Nucleic Acids Research 37(3), 858–865 (2009). doi:10.1093/nar/gkn1006

41. Altschul, S.F., Gish, W., Miller, W., Myers, E.W., Lipman, D.J.: Basic Local Alignment Search Tool. Journal of Molecular Biology 215(3), 403–410 (1990). doi:10.1016/S0022-2836(05)80360-2

42. Gusfield, D., Balasubramanian, K., Naor, D.: Parametric Optimization of Sequence Alignment. Algorithmica 12(4), 312–326 (1994). doi:10.1007/bf01185430

43. Frith, M., Hamada, M., Horton, P.: Parameters for Accurate Genome Alignment. BMC Bioinformatics 11(1), 80 (2010). doi:10.1186/1471-2105-11-80

44. Cao, M.D., Dix, T.I., Allison, L.: A genome alignment algorithm based on compression. BMC Bioinformatics 11(1), 599 (2010). doi:10.1186/1471-2105-11-599

45. Allison, L., Wallace, C.S., Yee, C.N.: Finite-state models in the alignment of macromolecules. Journal of Molecular Evolution 35(1), 77–89 (1992). doi:10.1007/BF00160262

46. Solomonoff, R.: A Formal Theory of Inductive Inference. Information and Control 7(2), 1–22224254 (1964)

47. Wallace, C.S., Boulton, D.M.: An Information Measure for Classification. Computer Journal 11(2), 185–194 (1968)

48. Henikoff, S., Henikoff, J.G.: Amino Acid Substitution Matrices from Protein Blocks. Proceedings of the National Academy of Sciences 89(22), 10915–10919 (1992)

49. Cao, M.D., Dix, T.I., Allison, L., Mears, C.: A Simple Statistical Algorithm for Biological Sequence Compression. In: Data Compression Conference, pp. 43–52. IEEE, Utah (2007). doi:10.1109/DCC.2007.7. http://doi.ieeecomputersociety.org/10.1109/DCC.2007.7 http://ieeexplore.ieee.org/lpdocs/epic03/wrapper.htm?arnumber=41487

50. Cao, M.D., Dix, T.I., Allison, L.: A Biological Compression Model and Its Applications. In: Arabnia, H.R.R., Tran, Q.-N. (eds.) Software Tools and Algorithms for Biological Systems. Advances in Experimental Medicine and Biology, vol. 696, pp. 657–666. Springer, ??? (2011). doi:10.1007/978-1-4419-7046-6_67. http://www.springerlink.com/content/k472251142228850/

51. Cao, M.D., Dix, T.I., Allison, L.: Computing Substitution Matrices for Genomic Comparative Analysis. In: Theeramunkong, T., Kijsirikul, B., Cercone, N., Ho, T.-B. (eds.) Advances in Knowledge Discovery and Data Mining. Lecture Notes in Computer Science, vol. 5476, pp. 647–655. Springer, ??? (2009). doi:10.1007/978-3-642-01307-2_64. http://link.springer.com/10.1007/978-3-642-01307-2_64

